# Dynamics of viral index infections in novel hosts

**DOI:** 10.1101/2020.04.09.033928

**Authors:** Nardus Mollentze, Daniel G. Streicker, Pablo R. Murcia, Katie Hampson, Roman Biek

## Abstract

Whether a pathogen entering a new host species results in a single infection or in onward transmission, and potentially an outbreak, depends upon the progression of infection in the index case. Although index infections are rarely observable in nature, experimental inoculations of pathogens into novel host species have a long history in biomedical research. This provides a rich and largely unexploited data source for meta-analyses to identify the host and pathogen determinants of variability in infection outcomes. Here, we analysed the progressions of 514 experimental cross-species inoculations of rabies virus, a widespread zoonotic pathogen which in nature exhibits both dead end infections and varying levels of sustained transmission in novel hosts. Inoculations originating from bats rather than carnivores, and from warmer to cooler-bodied species caused infections with shorter incubation periods that were associated with diminished virus excretion. Inoculations between distantly related hosts tended to result in shorter clinical disease periods, which will also impede transmission. All effects were modulated by infection dose and together suggest that increased virulence as host species become more dissimilar is the limiting factor preventing onward transmission. These results explain observed constraints on rabies virus host shifts, allow us to evaluate the risk of novel reservoirs establishing, and give mechanistic insights into why host shifts are less likely between genetically distant species. More generally, our study highlights meta-analyses of experimental infections as a tractable approach to quantify the complex interactions between virus, reservoir, and novel host that shape the outcome of cross-species transmission.

**Significance statement:** Emerging disease epidemics often result from a pathogen establishing transmission in a novel host species. However, most cross-species transmissions fail to establish in the newly infected species for reasons that remain poorly understood. Examining cross-species inoculations involving rabies, a widespread viral zoonosis, we show that mismatches in virulence, which are predictable from host and viral factors, make sustained transmission in the novel host less likely. In particular, disease progression was accelerated and virus excretion decreased when the reservoir and novel host were physiologically or genetically more dissimilar. These mechanistic insights help to explain and predict host shift events and highlight meta-analyses of existing experimental inoculation data as a powerful and generalisable approach for understanding the dynamics of index infections in novel species.

## Introduction

Cross-species transmission is an important source of emerging and endemic disease. Viruses such as West Nile virus, rabies virus and Lassa virus cause tens of thousands of human infections annually through transmission from animal reservoirs (Chancey et al., 2015; Hampson et al., 2015; Ogbu et al., 2007). Cross-species transmission is also the first step towards host shifts, where pathogens establish transmission cycles in novel hosts (Wolfe et al., 2007). While the broader-scale epidemiological and ecological factors driving cross-species transmission are beginning to be understood (reviewed in Lloyd-Smith et al., 2009; Plowright et al., 2014, 2017), we remain unable to anticipate whether cross-species transmission will cause ‘dead-end’ infections or lead to sustained transmission. Infection dynamics at the cross-species interface, specifically the probability of infection given exposure and the progression of index infections in novel hosts, are generally unobservable in nature. This is a crucial gap given that the outcomes of cross-species infections have profound implications for host shifts and disease emergence.

Cross-infection studies, in which viruses from a natural reservoir are experimentally inoculated into novel host species, provide a rare view into the dynamics of index infections. Since the dose, route, timing and origin of viral exposure are known, these factors can be controlled for to identify the ecological and evolutionary rules that govern the outcomes of cross-species transmission. We focus on *Rabies lyssavirus* (family *Rhabdoviridae*) as a model pathogen for understanding cross-species transmission (Fisher et al., 2018). Rabies virus is a zoonotic RNA virus, transmitted through an infectious bite, that infects all mammals and, untreated, has the highest case fatality ratio of any infectious disease (Hemachudha et al., 2002; Rupprecht et al., 2002). Rabies virus naturally infects multiple carnivore and bat species, which each perpetuate species-specific maintenance cycles (Mollentze et al., 2014). Although most cross-species transmission events do not lead to onward transmission, each maintenance cycle represents a rare past cross-species transmission event that established transmission in a novel host. Dead-end cross-species transmissions and historical host shifts are detectable in rabies virus phylogenies, and epidemiological surveillance reveals that nascent hosts shifts remain commonplace (Kuzmin et al., 2012; Mollentze et al., 2014; Streicker et al., 2010). As such, rabies virus exhibits extensive variation in the epidemiological outcomes of cross-species transmission. Here, we exploit cross-infection studies conducted over several decades, in which diverse mammalian species were inoculated with rabies viruses of bat and carnivore origin, to investigate the individual-level outcomes of index infections.

The potential for sustained onward transmission of rabies virus is likely to depend on the incubation period (from bite to the appearance of clinical signs) and the duration of clinical signs prior to death (here, the clinical period) of infected hosts. Longer incubation periods are associated with greater distribution of virus through the central nervous system (Fekadu et al., 1982), spread to a wider range of tissues (Davis et al., 2013), and higher virus titres in the salivary glands, all of which should facilitate onward transmission (Baer and Bales, 1967). Conversely, faster progression of infection has been associated with lower virus excretion, and in extreme cases animals die before the virus reaches the salivary glands, making transmission highly unlikely (Baer and Bales, 1967; Charlton et al., 1987; Davis et al., 2013; Fekadu et al., 1982). Further, the clinical period of rabies coincides with the period of greatest infectivity, when excretion of virus in the saliva often coincides with clinical signs such as aggression which promote transmission (Hanlon, 2013). Testing for shifts in incubation and clinical period durations, and in the amount of virus excreted, allows us to examine what constrains onward transmission of rabies in index hosts following cross-species transmission.

Based on previous work on rabies virus and in other host-pathogen systems, several mechanisms can be hypothesised to influence infection dynamics and thus the outcome of cross-species transmission:

1. Features of exposed host species (*host effects*) influencing the outcome of cross-species transmissions, irrespective of the infecting virus. For example, larger-bodied species may be more resistant to infection and thus require either higher infectious doses or a longer period of virus replication before symptoms become apparent. More generally, evolutionarily conserved similarities in host physiology mean that groups of related taxa might have similar susceptibility or clinical outcomes of infection (Longdon et al., 2011).
2. Features inherent to the virus lineage involved (*virus effects*), irrespective of the infected host, likely due to adaptation of individual lineages to reservoir host species. For example, key differences in disease expression between rabies viruses adapted to bats and those adapted to carnivores have been noted in humans, although it remains unclear whether this is a feature of the virus or due to differing routes of exposure (Begeman et al., 2018). Ultimately, the effect of a specific virus and its reservoir host cannot be disentangled.
3. *Host-virus interactions.* Both initial cross-species transmission and successful establishment occur most often between closely related hosts, often referred to as the phylogenetic distance effect (Gilbert and Webb, 2007; Longdon et al., 2015, 2011; Streicker et al., 2010). However, the mechanisms underlying this pattern remain obscure. Mammals also exhibit considerable variability in physiological features that are only moderately constrained by phylogenetic relatedness, such as body temperature (Clarke and Rothery, 2008). This may create distantly related pairs of reservoir and novel host species which nevertheless share key physiological features affecting disease outcome (Longdon et al., 2014), a potential explanation for the occurrence of host shifts over wide phylogenetic scales (e.g. the bat to primate host shifts involving SARS-coronavirus and Ebola viruses).

Here, we test these hypotheses by conducting a meta-analysis of individual-level data from 514 published experimental cross-species infections involving rabies virus. We show that features of the virus and of the inoculated host species interact with the initial conditions of exposure to influence the outcome of cross-species transmission in ways expected to affect the likelihood of onward transmission in the novel host species.

## Results

Our meta-analysis of cross-species inoculation experiments yielded results from 20 mammal species (in the orders Carnivora, Chiroptera, Cetartiodactyla and Rodentia), inoculated with 39 unique inocula from 7 reservoir species in the orders Carnivora and Chiroptera (Figure 1 & Table 1). Three outcome measures were recorded – the duration of the incubation period (N=443) and clinical period (N=175) and the amount of virus excreted (N=278).

**Table 1:** Summary of data available for each response variable

|  | Incubation<br>period | Clinical period | Presence of virus<br>in salivary glands* |
| --- | --- | --- | --- |
| Individual inoculations | 443 | 175 | 514 (278) |
| Studies | 19 | 14 | 23 (16) |
| Experiments | 25 | 19 | 30 (20) |
| Inocula | 35 | 17 | 39 (22) |
| Source taxonomic orders / species | 2 / 7 | 2 / 4 | 2 / 7 (2 / 4) |
| Inoculated taxonomic orders /<br>species | 4 / 19 | 4 / 13 | 4 / 20 (4 / 18) |
| Source-inoculated species<br>combinations | 30 | 19 | 33 (25) |

\*Joint fit, dataset includes inoculations providing data for incubation period only. Data for inoculations specifically providing information on the virus tires in salivary glands given in brackets

**Figure 1:**
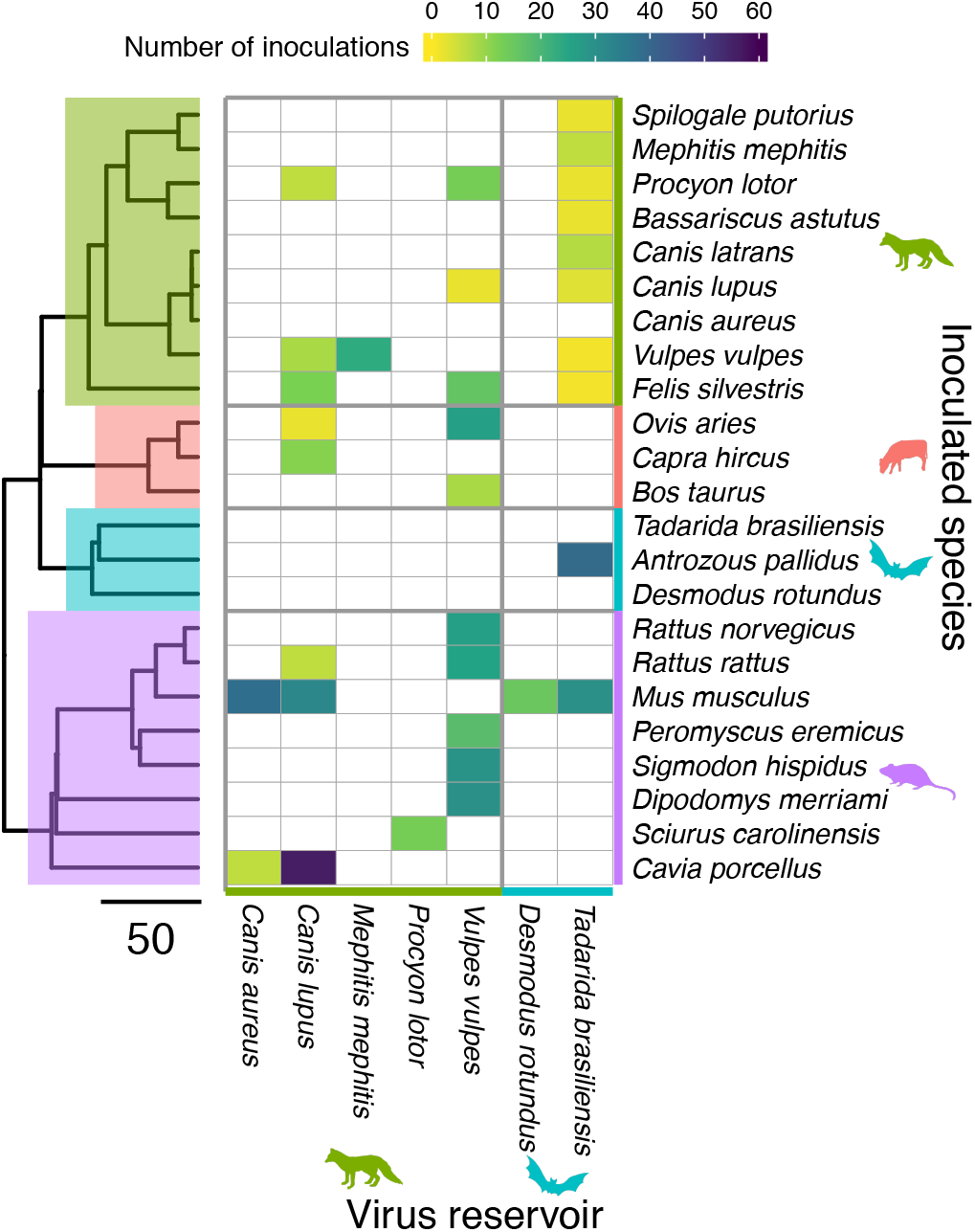
Source-inoculated species combinations for which data were included in this study, and the number of animals inoculated. Phylogenetic relationships are demonstrated using a composite time-scaled phylogeny, with Carnivora highlighted in green, Artiodactyla in red, Chiroptera in blue, and Rodentia in purple. Branch-lengths are in millions of years. The shading of cells indicates the number of inoculated animals for which data are available, with white cells indicating combinations for which no data were available. Three reservoirs occurred as sources only, receiving no inoculations which met our inclusion criteria (*C. aureus*, *T. brasiliensis* and *D. rotundus*). These species were nevertheless retained on the y-axis to demonstrate phylogenetic relationships.

### Incubation period

The time period between inoculation and the appearance of symptoms was highly variable, with a median duration of 15 days. While incubation periods ranged between 4 and 141 days, 95% lasted ≤ 28 days (all estimates based on a non-parametric Kaplan-Meier fit to the censored event times).

We modelled incubation period duration using log-normal generalized linear mixed models (GLMMs), correcting for phylogenetic non-independence among inoculated species and among reservoir species, as well as for clustering within experiments (see Methods). As expected, incubation periods were shortened by both increased dose and by inoculation routes which were closer to the brain, although the effect size estimate for inoculation distance included zero (Figure 2A). More importantly, the duration of incubation periods was also influenced by features of the virus as well as its interaction with the inoculated host. Specifically, significant differences in incubation period duration were associated with reservoir type (bat vs. carnivore) and with body temperature differences between source and inoculated hosts (Figure 2A). Both effects depended on viral dose. At low doses, viruses from bat reservoirs were associated with shorter incubation periods compared to viruses from carnivores, though this effect diminished at higher doses (Figure 2B). A similar effect was seen when the inoculated species was a known rabies reservoir (Figure 2C) although the 95% highest posterior density interval (HPD) of this effect size included zero (Figure 2A). The difference in typical body temperature between the virus reservoir and the inoculated species had a more marked effect on incubation period durations (Figure 2A). The onset of symptoms was delayed when the virus was inoculated into species with a warmer body temperature than its reservoir (negative values in Figure 2D), although this delay reduced with higher doses. The opposite was also true – in hosts with lower body temperatures than the virus reservoir, the incubation period tended to be shorter (Figure 2D, Supplementary figure S 1). Models fitting effects for inoculated and reservoir species body temperature separately allowed us to explore this temperature effect further. Inoculated species with higher typical body temperatures tended to have longer incubation periods (Supplementary figure S 2). However, viruses from reservoirs with higher body temperatures were associated with shorter incubation periods, suggesting that these viruses had adapted to counteract any losses in efficiency caused by the body temperature of their reservoir host. Importantly, there was no correlation between phylogenetic distance and body temperature difference among species (Supplementary figure S 3), indicating that the observed temperature effects were not explainable by the level of taxonomic relatedness among species. All fixed effects combined explained 19.2% of the variation in incubation period durations (HPD: 2.8 – 40.2%).

**Figure 2:**
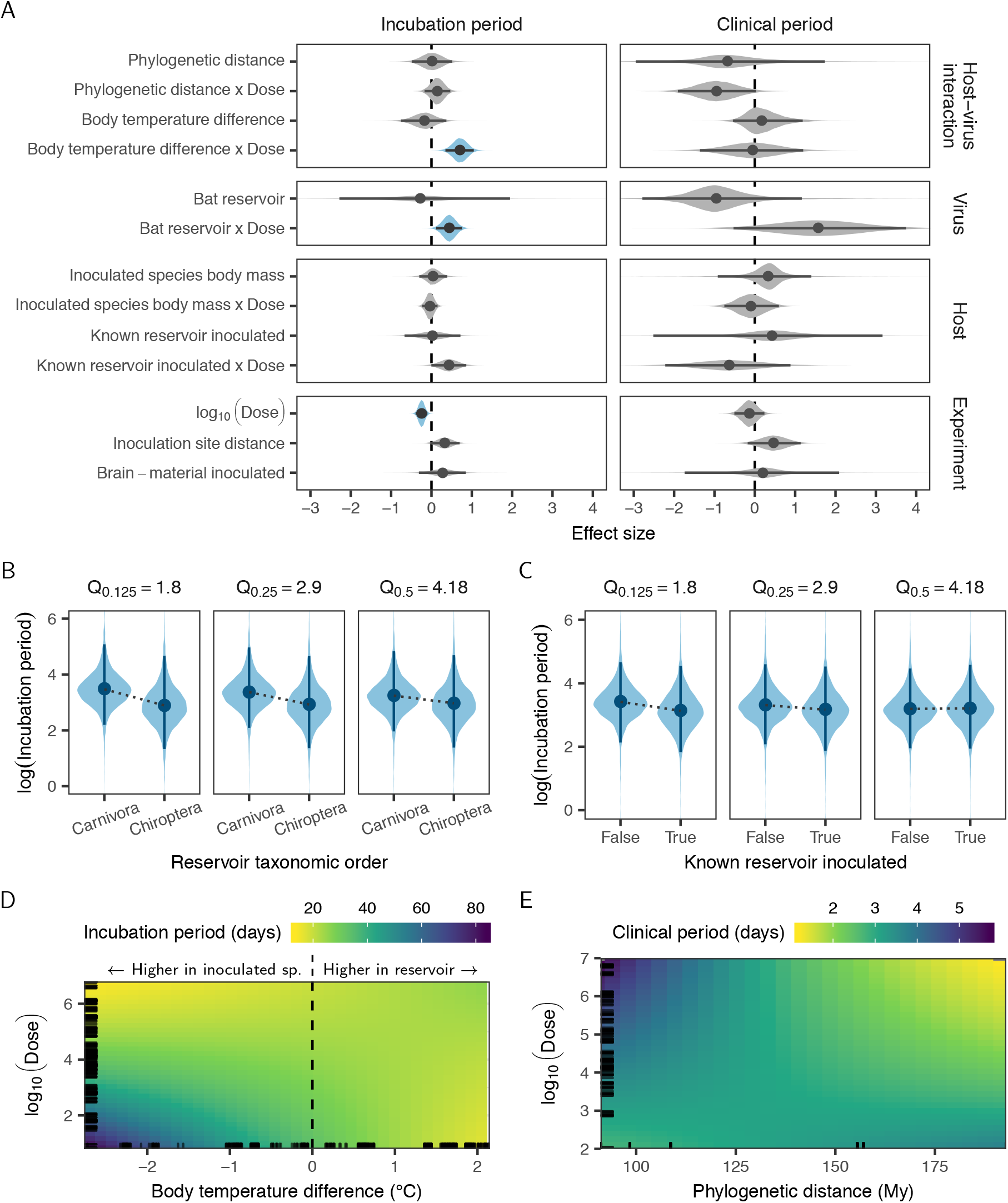
Factors affecting the progression of disease following cross-species inoculations. A: Coefficient estimates from independent models fit to the duration of incubation and clinical periods. In addition to the variables shown, both models contained corrections for non-independence between observations from the same experiment and for inoculated species phylogeny. The regression on incubation period durations additionally contained corrections for reservoir phylogeny. Lines indicate the extent of the 95% highest posterior density (HPD), while points indicate the posterior median. Shaded areas show the posterior distribution, with colour used to indicate estimates whose 95% HPD excludes zero. B-D: Predicted incubation period durations when varying dose along with reservoir taxonomic order (B), whether or not the inoculated species was a known reservoir of rabies virus (C), or the difference in typical body temperatures between the virus reservoir and the inoculated species (specifically reservoir temperature minus inoculated species temperature, D). For each set of predictions, all other explanatory variables in the model were held constant at their median observed value. In B & C, each sub-panel shows the predicted effect at different quantiles (Q) of observed doses (in log_10_ mouse LD50), indicated above the panel. E: Predicted duration of clinical periods as a function of phylogenetic distance and dose. Panels D-E show the posterior median of predictions.

### Clinical period

Once symptoms appeared, the median time to death (the clinical period) was 3 days, ranging from < 1 day to 8 days. We modelled clinical period duration using log-normal GLMMs, correcting for phylogenetic clustering among inoculated species and for clustering within experiments. In contrast to results for the incubation period, phylogenetic distance between reservoir and inoculated host species appeared to affect the duration of clinical periods. Cross-species inoculations between phylogenetically more distant species were associated with an increased sensitivity to high viral doses, resulting in shorter clinical periods (Figure 2E). However, the overall effect size estimate for this interaction included zero (Figure 2A). Bat-associated viruses appeared to have shorter clinical periods, and - as also observed for incubation periods - this effect depended on dose, but here it was poorly estimated, with the HPD again including zero (Figure 2A). All other fixed effects were small, and none could be clearly separated from zero, but combined the fixed effects explained 40.3% of the variation in clinical period duration (HPD: 12.9 - 64.0%).

### Virus titre in salivary glands

Onward transmission of rabies virus, which is mediated by an animal bite, requires presence of the virus in sufficiently high titres in the salivary glands. To test how the host-virus context of cross-species transmission affects the amount of virus excreted, we investigated the virus titre detected in salivary glands *post mortem* as a proxy. To simultaneously investigate potential explanations for the previously reported correlation between salivary gland virus titre and incubation period duration (Baer and Bales, 1967; Davis et al., 2013; Fekadu et al., 1982), we modelled the virus titre excreted jointly with incubation periods using a multi-response log-normal GLMM. When accounting only for clustering within experiments, salivary gland titres showed a moderate positive correlation with incubation period duration (Pearson correlation: 0.298, 95% HPD: 0.115 - 0.471; Figure 3A). Thus, consistent with previous work, animals which experienced longer incubation periods tended to have more virus in their salivary glands *post mortem*. Part of this correlation is accounted for by the inoculated species phylogeny (Figure 3B). The remaining residual correlation is explained by differences in dose (Figure 3C), with higher doses leading to decreased salivary gland titres (Figure 3D). The inoculation of species with a lower body temperature than the reservoir also tended to reduce the virus titre in the salivary glands, although the size of this effect could not be estimated precisely enough to make it distinct from zero (positive body temperature differences, Figure 3D & E). A clearer effect was observed for the interaction of reservoir status and dose: at low doses, known rabies reservoir species produced higher virus titres in the salivary glands than non-reservoirs (Figure 3G). At very high doses, however, we detected no difference in salivary gland titres, possibly because animals succumb too fast for any differences to develop.

**Figure 3:**
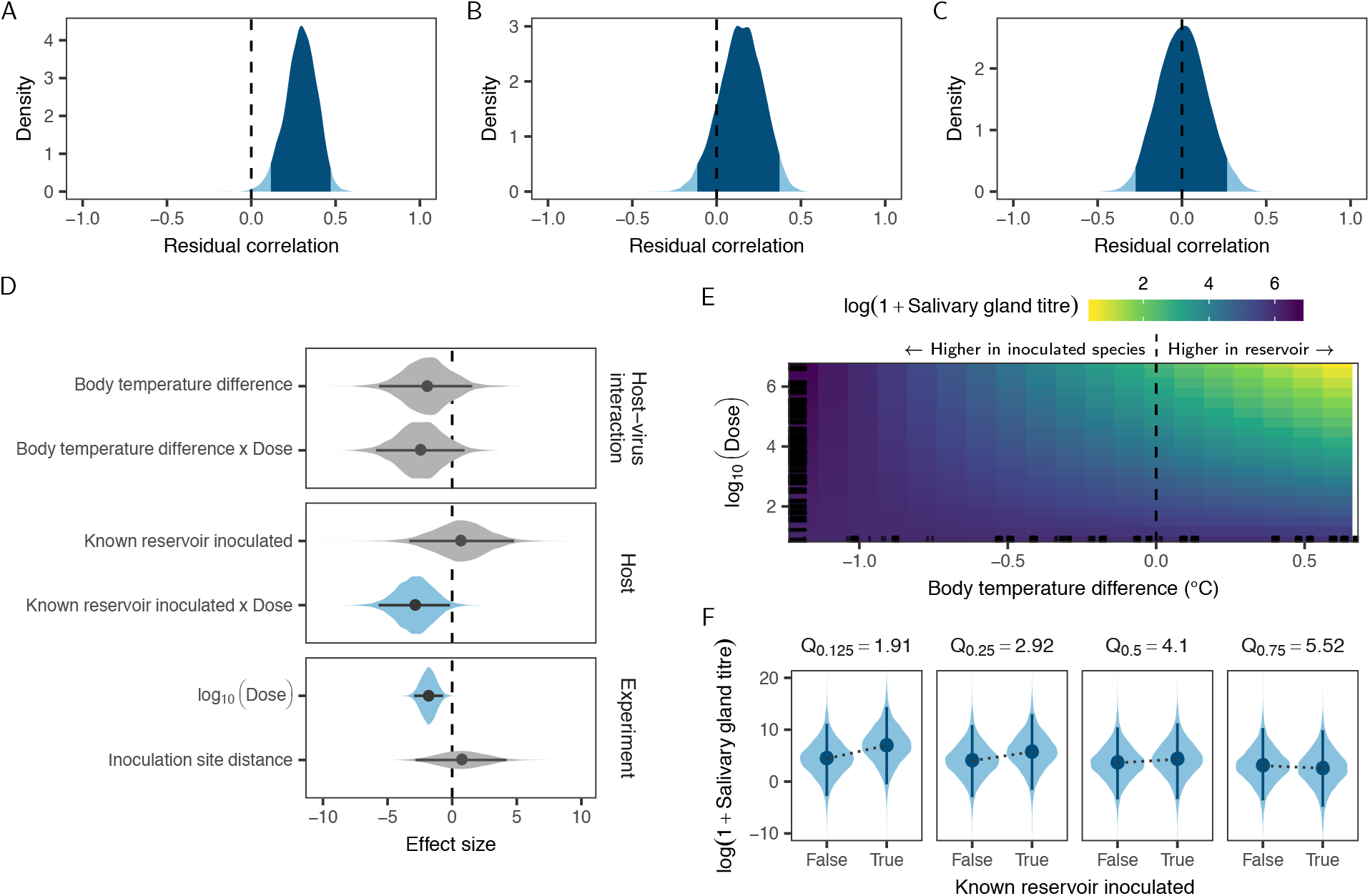
Factors predicting the titre of virus in salivary glands following cross-species inoculation. A: Correlation between virus titre and the duration of incubation periods, after accounting for clustering within experiments. Shown is the estimated posterior distribution from a multi-response regression fitted using the mcmcGLMM library in R, with the darker shaded area indicating the extent of the 95% highest posterior density (HPD). B-C: The remaining correlation is reduced when also accounting for correlation of virus titre and incubation period duration within the inoculated species phylogeny (B), and further reduced to approximately 0 when additionally accounting for differences in inoculated dose (C). D: Coefficient estimates for the regression on log virus titres in the salivary glands, when accounting for clustering within experiments and the inoculated species phylogeny. Lines indicate the extent of the 95% HPD, while points show the posterior median and shaded areas the shape of the posterior distribution, with colour used to indicate estimates whose 95% HPD excludes zero. E-F: Predicted salivary gland titres when varying dose and either the difference in body temperatures between the virus reservoir and the inoculated species (E) or whether or not the inoculated species is a known rabies virus reservoir (F), while keeping all other variables in the model constant. Panel E shows the posterior median of predicted virus titres. In F, predictions are shown at different quantiles of dose (in log_10_ mouse LD50), as indicated above each sub-panel.

## Discussion

The progression of viral infections within the index host following cross-species transmission is a crucial determinant of onward transmission, but is generally unobservable in nature. By analysing a unique dataset of experimental cross-species infections, we demonstrate that phylogenetic distance and specific physiological differences between the host species involved alter the progression of infections in ways that are expected to influence whether transmission in the novel host is sustained.

The association between incubation period duration, the extent of centrifugal spread to other tissues, and the amount of virus excreted suggests a direct mechanism linking longer incubation periods to onward transmission. Species with higher body temperatures than the reservoir host tended to have longer incubation periods, specifically at lower viral inoculation doses (Fig. 2D). Although one might expect body temperature to be a phylogenetically conserved trait, we found no correlation with phylogenetic distance (Supplementary figure S 3A), and others have shown that body temperatures are clustered primarily at higher taxonomic levels (Clarke and Rothery, 2008). As a result, most bat species have lower body temperatures than most carnivore species, but the distributions of body temperatures across bat and carnivore species still overlap (Supplementary figure S 3B). The finding that incubation period duration is influenced by body temperature is consistent with *in vitro* experiments showing that temperature can affect the infectivity of rabies virus, possibly by altering the rate of cell-to-cell spread (Morimoto et al., 1996). However, the specific mechanisms that could shorten incubation periods in a novel host environment which is colder than the host environment to which the virus is adapted, remain to be identified.

Crucially, we found clear evidence for virus adaptation to host body temperature (Supplementary figure S 2), consistent with infection progression being matched to each host species. Given this host adaptation and relatively poor correspondence between body temperature differences and phylogenetic distance, the observed temperature effect may help explain rabies virus host shifts across large phylogenetic distances. For example, despite host shifts from bats into carnivores being generally very rare, rabies virus has shifted repeatedly from big brown bats (*Eptesicus fuscus*, 36 °C) to striped skunks, (*Mephitis mephitis*, 36.45 °C; Kuzmin et al., 2012; Tacutu et al., 2013). More generally, our results suggest that transmissions to species with warmer typical body temperatures than the current reservoir are more likely to become established, since this would be expected to result in longer incubation periods and higher virus excretion. This might explain observations suggesting sustained transmission of rabies virus lineages associated with common vampire bats (*Desmodus rotundus*, 35 °C) in sympatric frugivorous bats (*Artibeus lituratus*; 37.3 °C; Kobayashi et al., 2007; Obregón-Morales et al., 2017; Tacutu et al., 2013).

The observation that low doses of viruses from bat reservoirs resulted in shorter incubation periods relative to those from carnivores (Figure 2A & B) suggests increased infectivity and/or faster within-host spread in bat-adapted rabies viruses. Since our data were limited to viruses from two bat reservoirs - with 82% of these associated with one species, *Tadarida brasiliensis* (Figure 1) - it remains unclear whether this is a general feature of bat-associated rabies viruses. But, similar results have been observed in humans, where both incubation and clinical periods were shorter when the virus originated from bats rather than carnivores (Begeman et al., 2018). This bat-associated effect may be the result of body temperature differences - at 35 and 36 °C, the two bat reservoirs included had cooler body temperatures than almost all inoculated species (Supplementary figure 3). Alternatively, since bats are considerably smaller than known carnivore reservoirs and likely transfer much smaller volumes of saliva during transmission, bat-associated rabies viruses may be adapted to transmit at lower doses. Although it may be expected that smaller animals require less virus to become infected and that body mass would modulate subsequent disease progression, we found no evidence that the body mass of inoculated species or its interaction with dose affects the duration of incubation or clinical periods (Figure 2A).

Following the incubation period, the appearance of clinical signs of disease typically coincides with viral excretion and transmission. The duration of clinical signs is therefore crucial in determining whether an index host can transmit to conspecifics. Notably in the case of rabies virus, the clinical period coincides with the onset of signs such as aggression that facilitate onward spread through biting. It is also relatively short and invariably ends in death of the infected host, terminating transmission opportunities (Hanlon, 2013). Since many animals in our dataset were euthanised at some point after symptoms appeared (meaning the duration of their clinical period was right-censored) it was difficult to estimate effect sizes for the clinical period with enough precision to clearly distinguish them from zero (Figure 2A). However, there was some evidence that increased phylogenetic distances between virus reservoirs and inoculated species reduced the duration of clinical periods. Such increased virulence would mean that onward transmission becomes increasingly unlikely following cross-species transmission between more distant relatives. This is consistent with previous work showing that the number of successful rabies virus host shifts among North American bats decreases with phylogenetic distance (Streicker et al., 2010). This effect of phylogenetic distance on virulence and infectious period also provides further evidence that rabies virus adapts to specific reservoir species by altering disease progression.

Because rabies virus is generally transmitted via bite, the amount of virus excreted in the salivary glands will affect the probability of transmission from the index case. Further, the overall strong effects of dose we observed suggest that the amount of virus transferred to secondary cases will be a primary determinant of disease progression in secondary cases and hence further transmission in the new host population. Several authors have noted a positive correlation between the length of rabies virus incubation periods and subsequent virus excretion levels (Baer and Bales, 1967; Davis et al., 2013; Fekadu et al., 1982). Our results confirm this correlation across a wide range of host species combinations, and show that it can be partially explained by inoculation doses, with higher doses leading to both shorter incubation periods and decreased virus excretion. The remaining correlation is explained by clustering of disease progression kinetics on the inoculated species phylogeny, with related species having similar incubation periods and excreting similar amounts of virus (Figure 3A-C). After removing these sources of correlation, non-reservoirs tended to have lower virus titres in their salivary glands than established rabies virus reservoirs (Figure 3D & F), which may explain why the virus remains restricted to a relatively small number of reservoir hosts despite frequent spillovers to other species (Mollentze et al., 2014). Our results also provide tentative evidence of shorter incubation periods in non-reservoirs (Figure 2A). The exact mechanisms underlying these differences between known rabies virus reservoirs and other species remain unexplained, but are likely to be evolutionarily conserved, given the phylogenetic clustering in excretion levels apparent in our analysis.

Many of our results point to adaptation of the rate of disease progression to match individual host species. Studies of rabies virus host shifts have thus far failed to find sites in the virus genome which consistently change during host adaptation (Kuzmin et al., 2012; Streicker et al., 2012; Troupin et al., 2016). This has led to the suggestion that host adaptation can be achieved through numerous sets of molecular changes (Streicker et al., 2012), which would indeed be the case if the requirement is to balance disease progression to the point where onward transmission becomes likely. In some host-pathogen systems, such adaptation is explained by a trade-off between selection for faster growth to maximize viral load and thereby infectiousness, and selection for reduced host damage (virulence) to maximize transmission opportunities (de Roode et al., 2008; Fraser et al., 2007). In contrast, we observed a positive correlation between salivary gland titres and incubation period duration, implying that slower host damage leads to greater infectiousness. This, in turn, suggests a trade-off between faster replication and/or spread (speeding up disease progression) and the ability to reach the salivary glands via functioning neural pathways. Such a trade-off is supported by the observation that the correlation between incubation period duration and salivary gland virus titre was modulated in part by dose (Figure 3C), with higher amounts of virus reducing both.

The experimental data analysed here offer a unique view on index infection dynamics following cross-species transmission. By revealing the complex links between dose, physiological differences between hosts, disease progression and virus excretion, our analyses bring us closer to being able to model and predict the process of disease emergence and host shifts. The large dataset of controlled infections further enabled us to generate disease progression parameter distributions for all observed combinations of within and cross-species transmissions which can be directly applied in future efforts to model rabies transmission dynamics (supplemental dataset S1). Of note - many effects observed here were eventually overcome by high doses (Figure 2B-D & Figure 3F; the median dose for experiments was 40,000 mouse LD50s), and future infection studies should aim to utilize doses closer to those of natural exposures. Following cross-species transmission, rabies virus shows increased virulence (i.e., more rapid death) in more distantly related species, to the point that opportunities for transmission are likely to be markedly reduced. At the same time, a mismatch in host physiological features (including features not strongly correlated with phylogeny, such as body temperature) can alter both infectivity and disease progression, with implications for onward transmission. Thus, the picture that emerges is one of a potential virulence mismatch in index infections, that may explain why - despite having the ability to infect all mammals and frequent involvement in cross-species transmission events - rabies virus remains restricted to a relatively small number of species-specific maintenance cycles.

While the determinants of cross-species transmission have been the subject of intense research (reviewed in Lloyd-Smith et al., 2009; Plowright et al., 2014, 2017), the very next step, i.e. what happens during the initial infection to determine the likelihood of onward transmission, has remained relatively unexplored. Our results show that meta-analyses of cross-species infection experiments provide a tractable means of investigating this process. Expanding such analyses to other viruses may allow us to identify general rules which predict the outcome of cross-species transmissions. More work is needed to understand the host features that affect the probability of infection upon exposure, the within-host mechanisms driving virulence, and the epidemiological consequences of differences in disease progression and virus excretion. Our findings illustrate how understanding these mechanisms will be key to predicting which cross-species exposures are most likely to lead to future host shifts of rabies virus, and of zoonotic diseases more broadly.

## Methods

### Literature search and data collection

A search for published rabies virus infection studies was performed as described in supplementary document S1. Searching across the PubMed and Web of Science databases yielded 2279 records on 16 January 2015. These records were reviewed according to the criteria listed in table 1 to select studies for inclusion in the meta-analysis.

From each study, we recorded individual-level data on the species inoculated, the dose, inoculation route, and the reservoir host species of the virus used. Response variables, when available, included the observed incubation and clinical period durations, and the titre of virus present in the salivary glands post-mortem. Incubation and clinical period data comprised a mixture of exact times, interval censored times (i.e. studies only reported ranges for groups of animals), and right-censored observations (i.e. deaths unrelated to rabies before the conclusion of the study or euthanasia of survivors at the end of each study). Because animals are generally euthanised once symptoms appeared, the majority of data on the duration of clinical periods were right-censored, with exact durations reported in only a small number of studies, generally performed before euthanasia became common practice in animal experiments.

Taxonomic classifications were updated to match Wilson and Reeder (2005), by matching the scientific and common names given in each publication against the Integrated Taxonomic Information System database^1^. Data from our meta-analysis were supplemented with species-level data from the PanTHERIA database, along with body-temperature data from the AnAge database (Jones et al., 2009; Tacutu et al., 2013). Because not all records were resolved to sub-species level, and external data sources only contained data at the species level, information on the specific subspecies involved was ignored in the analyses described here. This resulted in two pairs of subspecies being clustered together, while two domesticated species were analysed using species-level data for their wild ancestor (Supplementary table S 1). Further data cleaning and validation steps are described in supplementary document S1.

### Accelerated failure time model

The durations of incubation and clinical periods were modelled using independent generalized linear mixed models on the censored event times, in this context more frequently termed accelerated failure time models. These models perform a regression on the waiting time to some specific event (e.g. the appearance of clinical signs, signifying the end of the incubation period), with coefficients acting to increase or decrease the time to the event. We assumed a log-normal distribution for the event times **T**.

Thus, the duration of the incubation period of each individual *i* of species *r* (the inoculated species), inoculated with a virus from species *d* (the source or reservoir species) in experiment *j* was modelled as:

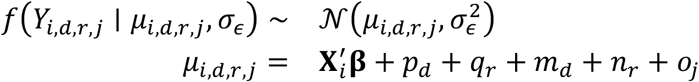

where **Y** = log(**T**), while *μ*_*i,d,r,j*_ and *σ*_*ϵ*_ are the mean and standard deviation of a normal distribution, respectively. Coefficients are represented by **β**_*i*_, with 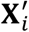 representing a vector of data on potential explanatory variables. Finally, (**p**, **q**) and (**m**, **n**, **o**) respectively represent phylogenetic and non-phylogenetic random effects for the source species (the virus reservoir, **p** and **m**), inoculated species (**q** and **n**), and experiment (**o**).

Phylogenetic random effects were drawn from a multivariate normal distribution taking the form

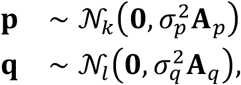

where **0** is a vector of zeros (of length *k* or *l*, equal to the number of source or inoculated species, respectively), and 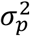 and 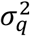 are variance parameters. **A**_*p*_ and **A**_*q*_ represent correlation matrixes for all source and inoculated species, respectively. These matrixes were calculated from a composite time-scaled phylogeny generated by timetree.org (Kumar et al., 2017) assuming a Brownian model of trait evolution using version 3.5 of the APE package in R (Hadfield and Nakagawa, 2010; Paradis et al., 2004). These random effects adjust for potential correlation in the response variables due to relatedness. Similar results were obtained when using the mammalian supertree of Bininda-Emonds et al. (2007), but this supertree had a slightly lower resolution than the timetree.org phylogeny.

The non-phylogenetic random effects took the form

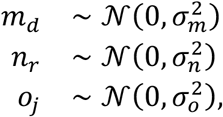

where 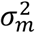 and 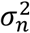 respectively measure the variance between source and inoculated species not captured by the Brownian model (Longdon et al., 2011), while 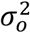 measures the variance between experiments. A similar model was used for the duration of clinical periods, except that data were pooled across virus reservoirs by removing the random effects for reservoir species and reservoir phylogeny (**p** and **m** above). This was necessary because the clinical period data involved viruses from just four reservoir species, making it impossible to accurately estimate the variance between observations associated with different reservoirs.

To accommodate censoring, the vector of event times, **T**, was treated as a latent variable. When only the range of incubation or clinical period durations was given for a specific group of animals, data was treated as interval censored, i.e. *T*_*i,d,r,j*_ ∈ [*L*_*i,d,r,j*_, *U*_*i,d,r,j*_], where **L** and **U** represent the lower and upper boundaries of observed event times (supplementary document S1). When an animal left the dataset without experiencing an event (e.g. when it was euthanised), the time of censoring was recorded as a lower boundary, reflecting our knowledge that the event of interest had not yet occurred by this point, but could have occurred at any time after this. In this case *T*_*i,d,r,j*_ ∈ [*L*_*i,d,r,j*_, ∞).

In this context, exact observations can be recorded simply by setting *L*_*i,d,r,j*_ = *U*_*i,d,r,j*_.

### Multi-response models

In an independent model, the amount of virus detected in salivary glands *post mortem* was modelled jointly with incubation period durations, to allow estimation of the amount of residual correlation between incubation period duration and the amount of virus in the salivary glands. Several authors have noted a link between these measures (Baer and Bales, 1967; Davis et al., 2013; Fekadu et al., 1982), but it remains unexplained.

This regression was similar to the model above, except that the normal distribution on log(observations) was replaced with a multivariate normal distribution. Thus, in the full model, the observed value *Y* of response variable *v* for individual *i* of species *r* in experiment *j* was modelled as

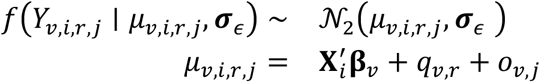

where *v* = 1 represents the incubation period (i.e. ***Y***_*v*=1_ = log (***T***)) and *v* = 2 is the virus titre in the salivary glands. Virus titre was thus also modelled as log-normal, i.e. ***Y***_*v*=2_ = log(1 + ***W***), where ***W*** represents the observed titres, which may be 0 if no virus was detected in the salivary glands). **β**_*v*_ is a vector of coefficients unique to each response variable, while q_*v,r*_ and o_*v,r*_ are random effects for inoculated species phylogeny and experiment, respectively.

In these models, ***σ***_*ε*_ is a variance-covariance matrix of the form

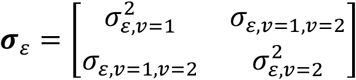

where 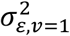 is the residual variance in response variable 1 (incubation period durations), and *σ*_*ε*,*v*=1,*v*=2_ is the residual covariance between incubation period durations and salivary gland titres. From this, the Pearson correlation between incubation periods and salivary gland titres can be calculated as

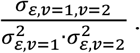

### Explanatory variables

Variables measuring differences between the reservoir and the inoculated species were included to assess the influence of previous virus adaptation on the outcome of infection in heterologous host species. These included the phylogenetic distance between the reservoir and the inoculated species, measured as patristic distances along the same composite time-scaled phylogeny generated by timetree.org used above. As above, similar results were obtained when using the mammalian supertree of Bininda-Emonds et al. (2007). We also included the difference in typical body temperatures between the reservoir and inoculated species, as an example of a physiological difference which does not appear to follow phylogenetic constraints (Clarke and Rothery, 2008), because temperature is known to affect rabies virus infectivity *in vitro* (Morimoto et al., 1996). Finally, a binary variable distinguishing viruses derived from bat and carnivore reservoirs was included, because differences in the clinical presentation of bat- and carnivore-associated rabies virus infection in humans have been noted (Begeman et al., 2018).

Features of the inoculated host species where accommodated primarily through random effects for species and inoculated species phylogeny. However, we also included a measure of the typical body mass of the inoculated species, since larger species may be proportionally more resistant to the effects of a given dose of virus. Because only some species maintain rabies virus transmission endemically, for reasons that are not well understood, a binary variable distinguishing known reservoirs of rabies virus from other inoculated species was also included.

Differences between experiments were accommodated by including variables for dose, the inoculation site, and whether the inoculum consisted of brain material or was derived from salivary glands/saliva, along with a random effect distinguishing between experiments to accommodate any remaining differences. Because the doses encountered in these experiments differed over several orders of magnitude and the effects of increasing dose is assumed to decrease (saturate) at very large doses, this variable was included in its log-transformed form. The varying inoculation routes encountered were summarised as a ‘proportional inoculation distance’, representing the relative distance between the inoculation site and the brain (the primary site of rabies virus replication). This distance was calculated by classifying inoculation routes by body part (head, neck, torso or limbs) and depth (intracranial, intramuscular, or subcutaneous) and was expressed as a proportion, where 1 indicates the furthest and shallowest possible inoculation site relative to the brain (sub-cutaneous inoculation of a limb), while 0 indicates intracerebral inoculation (Supplementary table S 2 & Supplementary table S 3). Such proportional scaling means this variable is independent of the differing body sizes of the inoculated species. Finally, because larger doses may compensate for any decreases in infectivity caused by features of the inoculated species and/or physiological differences between the inoculated species and the reservoir to which the virus was adapted, we included interactions between dose and all host and virus effects above.

### Model fitting

Models were fit using version 2.25 of the MCMCglmm package in R version 3.5.1 (Hadfield, 2010; R Core Team, 2018). All coefficients and the residual variance parameter received the default prior distributions used by MCMCglmm, while parameter-expanded priors were used for the variance parameters of all random effects (Gelman, 2006). For each dataset, models were fitted using 10 million MCMC steps, saving every 1000th sample. The first 10% of samples in each chain were discarded as burn-in. Results were inspected and summarised using version 0.18-1 of the coda package in R (Plummer et al., 2006). Effective sample sizes were checked to ensure efficient sampling was achieved, and chains were visually inspected for convergence.

## Acknowledgments

We thank Paul Johnson, Mafalda Viana, Dan Haydon and Ben Longdon for helpful discussions and advice on statistical analyses, and Andrew Yates for comments on a draft on this manuscript. NM was funded by a Lord Kelvin - Adam Smith studentship from the University of Glasgow. PRM was funded by the Medical Research Council (MC_UU_12014/9). DGS was supported by a Sir Henry Dale Fellowship, jointly funded by the Wellcome Trust and Royal Society (102507/Z/13/Z) and a Wellcome Senior Research Fellowship (217221/Z/19/Z). KH was supported by the Wellcome Trust (207569/Z/17/Z & 095787/Z/11/Z).

## Competing interests

The authors declare no competing interests.

## Supplementary material

### Supplementary tables & figures

**Supplementary table S 1:** Known sub-species in the dataset, which were analysed at the species level

| Species | Subspecies | Common name |
| --- | --- | --- |
| *Antrozous pallidus* | *pallidus* | Pallid bat |
| *Canis lupus* | *familiaris* | Domestic dog |
| *Mustela putorius* | *furo* | Domestic ferret |
| *Tadarida brasiliensis* | *mexicana* | Mexican free-tailed bat |
|  | *brasiliensis* | Brazilian free-tailed bat |
| *Vulpes vulpes* | *crucigera* | European red fox |
|  | *fulves* | Eastern North American red fox |

**Supplementary table S 2:** Classification of inoculation routes by body zone (rows) and depth (columns)*

|  | *Intracranial* | *Intramuscular* | *Subcutaneous/peripheral* |
| --- | --- | --- | --- |
| *Head* | Intracerebral, Optic nerve, Intracranial, Intracisternal, IC, IC intercranial | IM-masseter, IM-temporal, IM occipital, Submandibular, IM-masseter, Masseter, IM masseter, IM-Masseter, IM-masseter(both.sides), IM-masseter(both), left & right masseter, IM Right Masseter, IM M.masseter, IM Left Masseter, IM-maseter | Intranasal, Intranasally, Subcutaneous-lips |
| *Neck* |  | IM-neck, IM-cervical, IM-cervical(left), IM-deap-neck | Subcutaneous-neck, subcutaneous(neck) |
| *Torso* |  | Deltoid, IM-deltoid, IM-Deltoid, IM-pectoral, IM-Dorsal | SC-deltoid |
| *Limbs* |  | IM-quadriceps femoris (right), IM-right hindleg, IM-hind leg, IM-leg, IM right thigh, IM-LongDigitalExtensor(Pelvic), IM-AbductorDigitiQuinti(Pelvic), IM-gastrocnemius, Right gastrocnemius muscle, Gastrocnemicus , IM-crural, IM-HindFoot | Peripheral-footpad, SC-HindLimb |

\*The inoculation routes shown are free-text entries present in the raw data, where IC = intracranial, IM = intramuscular, and SC = subcutaneous.

**Supplementary table S 3:** Distances assigned to each inoculation route category to express relative distance to the brain*

|  | *Intracranial (0)* | *Intramuscular (1/7)* | *Subcutaneous (3/14)* |
| --- | --- | --- | --- |
| *Head (1/14)* | 0 | $1/14 + 1/7$ | $1/14 + 3/14$ |
| *Neck (1/7)* | - | $1/7 + 1/7$ | $1/7 + 3/14$ |
| *Torso (3/7)* | - | $3/7 + 1/7$ | $3/7 + 3/14$ |
| *Limbs (7/7)* | - | $7/7 + 1/7$ | $7/7 + 3/14$ |

\*Distances were chosen to ensure intramuscular inoculations to the neck received the same value as subcutaneous inoculations to the head, and were subsequently scaled to range between 0 (intracranial, head) and 1 (subcutaneous, head). See supplementary document S1 for details.

**Supplementary figure S 1:**
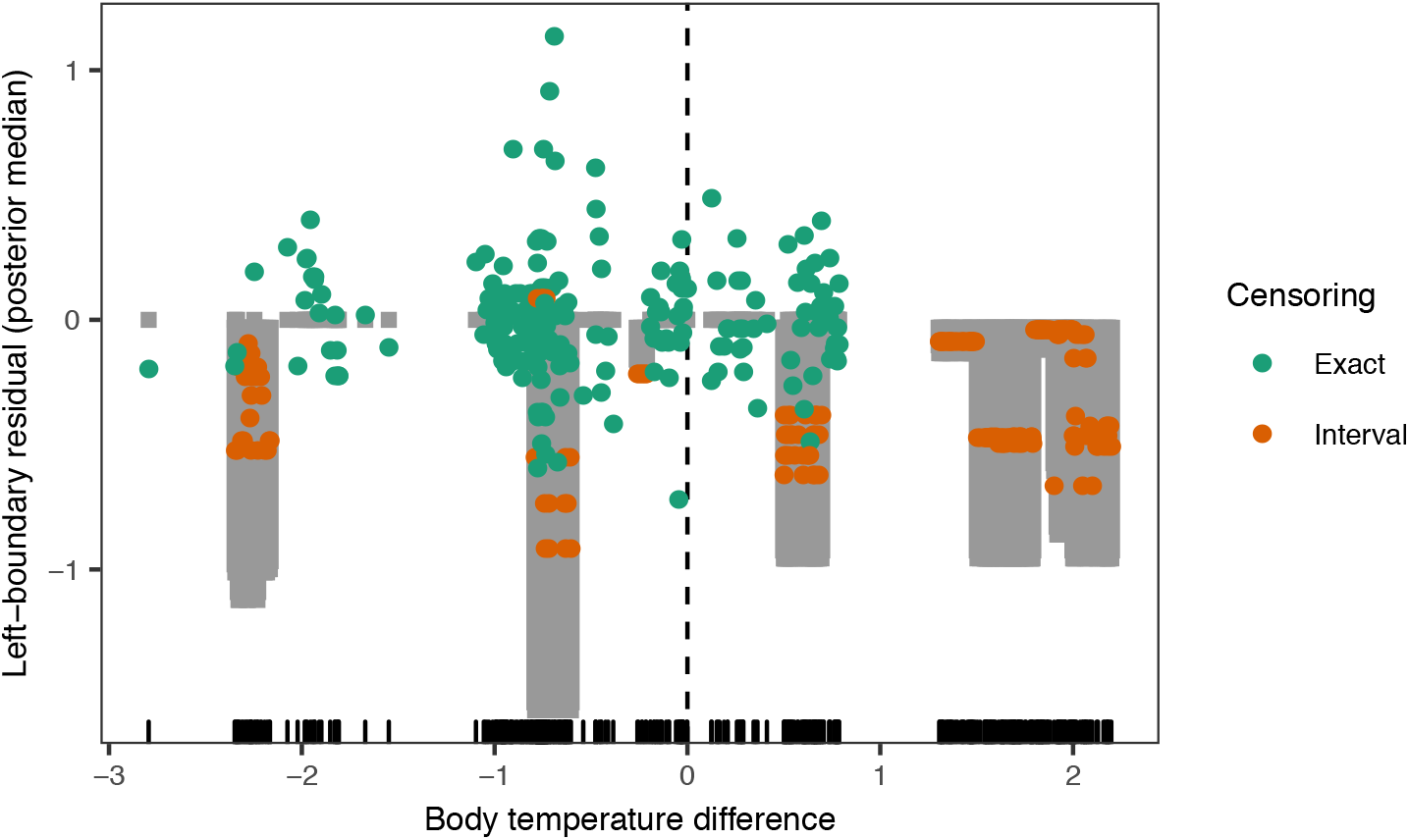
Posterior median residuals as a function of body temperature difference. The observed data was subject to censoring, but the lower-boundary was always known. Residuals were calculated as the observed lower limit minus the fitted value, with grey-shaded areas showing the expected range of each observation. In the case of exact observations, a perfect fit would result in residuals equal to 0, while the residuals for interval-censored observations are expected to lie in [0, lower - upper). No systematic bias in residuals was observed, supporting the conclusion that the effect of body temperature distance remained linear for both positive and negative values.

**Supplementary figure S 2:**
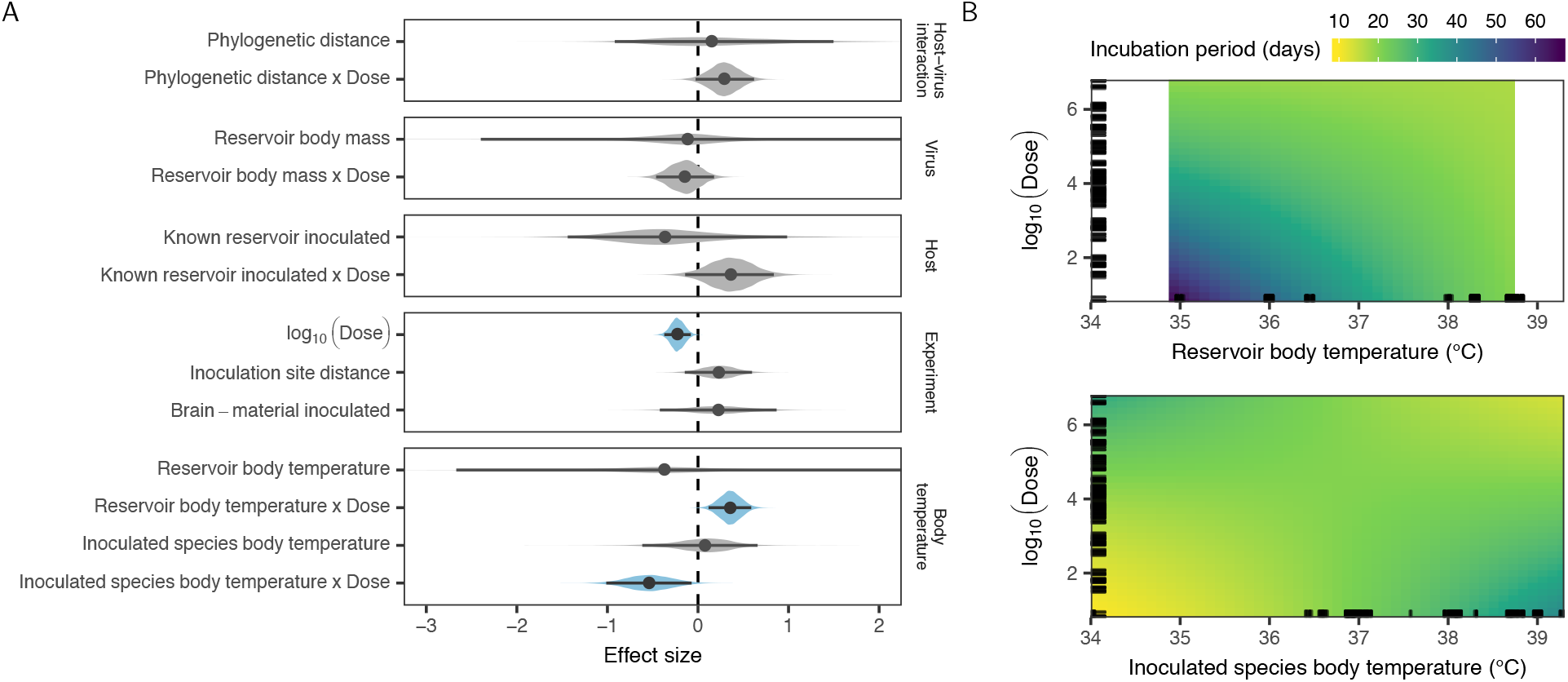
Coefficient estimates and predicted effects when fitting separate effects for the typical body temperature of the inoculated species and virus reservoir. A: Coefficient estimates, with lines showing the 95% highest posterior distribution and points showing the posterior median. Compared to the model shown in the main text, this model does not contain an effect for inoculated species body mass (which had no effect in the full model, and is correlated with inoculated species body temperature) and contains an effect for reservoir body mass, replacing the variable separating bat-associated viruses from carnivore-associated viruses (which was correlated with inoculated species phylogeny). B: Posterior median predicted incubation periods as a function of reservoir (top) or inoculated species body temperature (bottom) and dose. Predictions are shown within the range of observed values in the dataset.

**Supplementary figure S 3:**
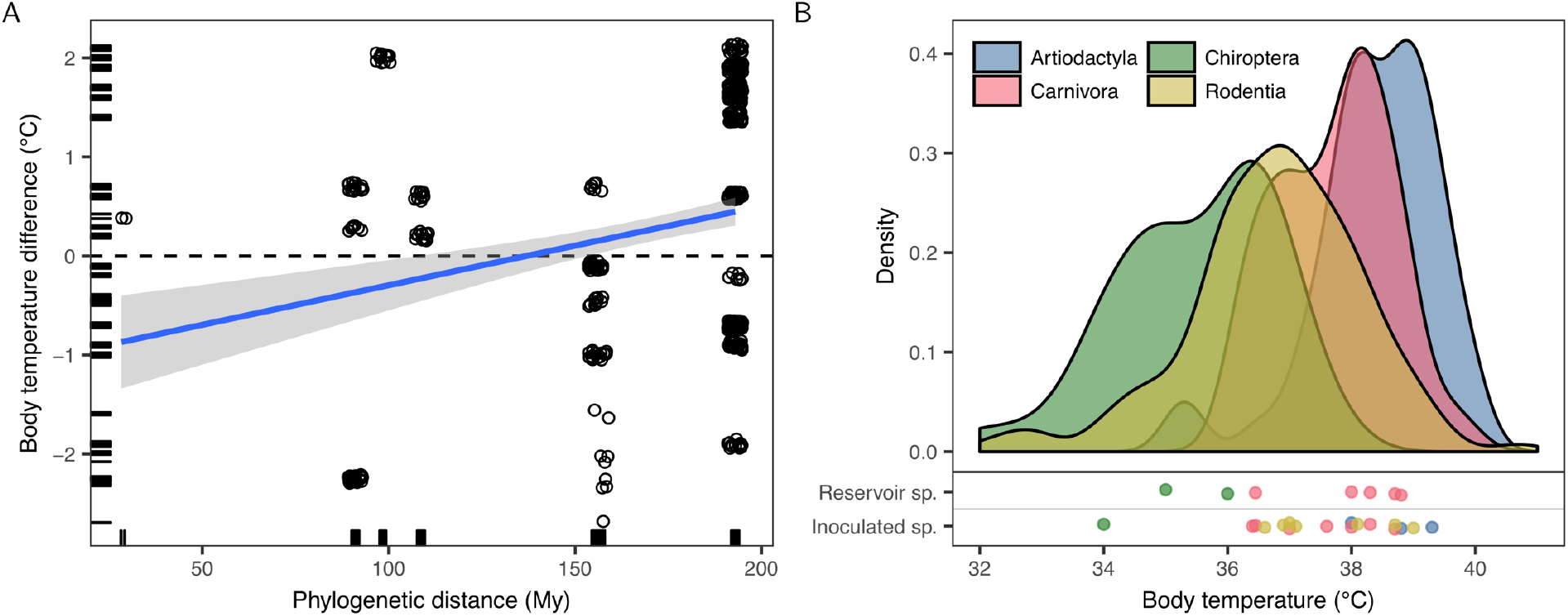
Relationship between phylogenetic distance and host body temperature. A: Relationship between phylogenetic distance (in millions of years) and body temperature difference in the incubation period dataset. Points are shown jittered for clarity, while the blue line represents the best linear fit. B: Distribution of body temperature in the taxonomic orders of hosts included in this study, based on all species for which body temperature data are available in the AnAge database. Points indicate the body temperatures for species included in the incubation period dataset as either reservoirs or inoculated species, jittered vertically to reduce overlap.

### Supplementary document S1: Supplementary methods

#### Literature search

To allow optimisation of the literature search strategy, a set of publications of known relevance (according to the abstract review criteria, *Supplementary table S 4*) was obtained by manual searching and from citations in several rabies textbooks. A script implementing the BioPython library in Python 3 was used to find a search query that returned the greatest possible number of these publications in a PubMed search, whilst excluding the most common sources of false positives - notably annual veterinary reports and vaccine studies. This resulted in the following query for Web of Science:

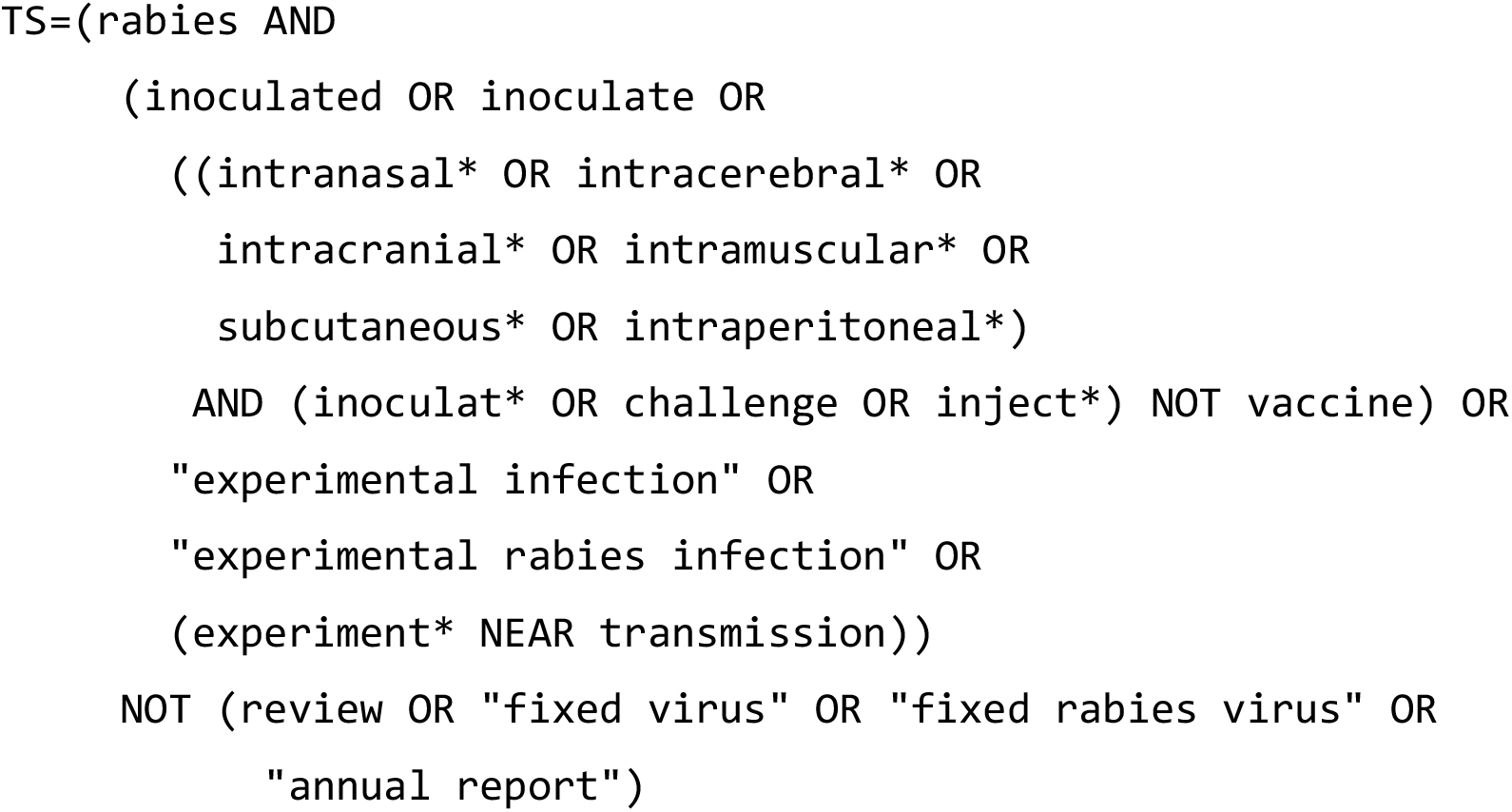

The same query was used to search the PubMed database, except that all instances of the NEAR operator were replaced with AND (PubMed does not support proximity searching). The PubMed version of this query returned 46 out of the 53 publications of known importance confirmed to be in the PubMed database. Of the remaining seven, five publications did not have searchable abstracts in the PubMed database.

Combining the results from Web of Science and PubMed yielded 2501 records on 16 January 2015. The search results were further filtered to exclude records where the keyword rabies did not occur in either the English title or abstract, although records where either of these fields was empty were also retained. The abstracts of the resulting 2279 records were reviewed according to the criteria in *Supplementary table S 4*. The abstracts of 35 records (1.54% of the filtered search results) could not be obtained in time for consideration. A total of 412 records (18.36% of the remaining search results) were selected for full text review, and the full texts of 382 publications (92.72%) were successfully obtained.

Publications in languages other than English were subjected to optical character recognition (where needed) using VietOCR version 4.0, a graphical user interface for the Tesseract OCR engine (version 3.30RC), optimised for each language being recognised. The digitised text was manually corrected and machine-translated using the statistical machine translation version of Google translate. All full texts were then reviewed using the criteria in *Supplementary table S 4*, resulting in the selection of 63 publications for inclusion in this study.

**Supplementary table S 4:** Criteria used to select publications

| Review stage | Criteria |
| --- | --- |
| Abstract review | Involves experimental inoculations with rabies virus<br>Does not solely focus on laboratory-adapted virus strain |
| Full text review | Source and inoculated species clearly identified<br>Source host species is a known maintenance host of rabies virus (passaging in non-maintenance hosts allowed)<br>Inoculated host species is a mammal<br>Inoculated host species is wild-type or outbred<br>Inoculated dose stated (in mouse LD<sub>50</sub>)<br>Healthy, unvaccinated animals used<br>Multiple experimental groups or a control group present |

#### Data cleaning and validation

Individual-level data on all inoculated animals present in the selected publications were recorded. Following data-entry, the raw data was extensively checked for accuracy and consistency. The scientific names of all inoculated and source species were manually checked for current validity using the ITIS database and standardised to match the taxonomy of Wilson and Reeder (2005). A further literature search was performed to obtain data on whether each of the included species is a known maintenance host of rabies virus. Records where the source species was not a known maintenance host were validated against the original publication to check for passaging and information on the maintenance host associated with the virus inoculum used. All virus inoculum identifiers were manually checked to identify the re-use of viruses in multiple publications, and were corrected to reflect this shared origin.

Some data were excluded from the current study, but retained in the database for potential future studies. A total of 197 records were excluded because the method of titration used to determine the dose inoculated was either not known (generally negative control animals) or because their reported inoculation dose involved a tissue culture-based rather than an *in vivo* method of titration. All remaining records listed virus doses obtained by intra-cerebral inoculation of mice to determine the 50% lethal dose. A further 31 records with a reported dose of 0, representing non-inoculated control animals, were also excluded. A total of 131 records involving five source species were excluded because the *source* species was either not a known maintenance host of rabies virus or could not be resolved to the species level, and 17 records involving one inoculated species were excluded because the *inoculated* species could not be resolved to species level.

A derived variable was created to unify diverse descriptions of inoculation routes and sites, based on the proportional distance of each inoculation site from the brain. Inoculation routes were classified into four zones on the body and three depths, and distances were assigned to each category (*Supplementary table S 2* & *Supplementary table S 3*). These distances were normalised to lie in [0,1]. Inoculation routes that were not specific enough to be classified into this scheme were checked against the original publications, and 115 records where the inoculation route information could not be improved were excluded.

All available data on the timing of disease progression and the total number of days survived following inoculation were recorded. When timing data was reported for groups of animals, the event times for each animal in the group was recorded as known only to be within the interval reported for that group (a form of interval censoring). Data on the total number of days survived was recorded in both exact (**T**_exact_) and interval censored (**T**_lower_ and **T**_upper_) forms, while the length of incubation and clinical periods were recorded in interval censored form only (with both columns given the same value when an exact time was available). Although only two of these timing variables needs to be known to calculate the third, data was recorded only in the form reported in each study.

The data were then processed as described in supplementary figures 12 - 13, and timing data were checked for internal consistency. This involved consolidating information on the cause of death, results of clinical observations and post-mortem diagnostic tests and the timing of disease progression stages associated with each animal into variables describing the lower and upper limits of the incubation and clinical period duration. When rabies was confirmed for a given animal through either clinical diagnosis or post-mortem diagnostic tests, it was assumed to have rabies. In many cases, data could be calculated for unreported timing variables by using some combination of reported data on either the number of days survived post inoculation, the duration of the incubation period, and/or the duration of the clinical period. For example, if only the earliest day that an animal *i* could have died (*T*_*i*,lower_) as well as the upper limit of the clinical period for that animal (*M*_*i*,upper_) is known, the lower limit of the incubation period can be calculated as *I*_*i*,lower_ = *T*_*i*,lower_ − *M*_*i*,upper_ (supplementary figure S12).

**Supplementary figure S 12:**
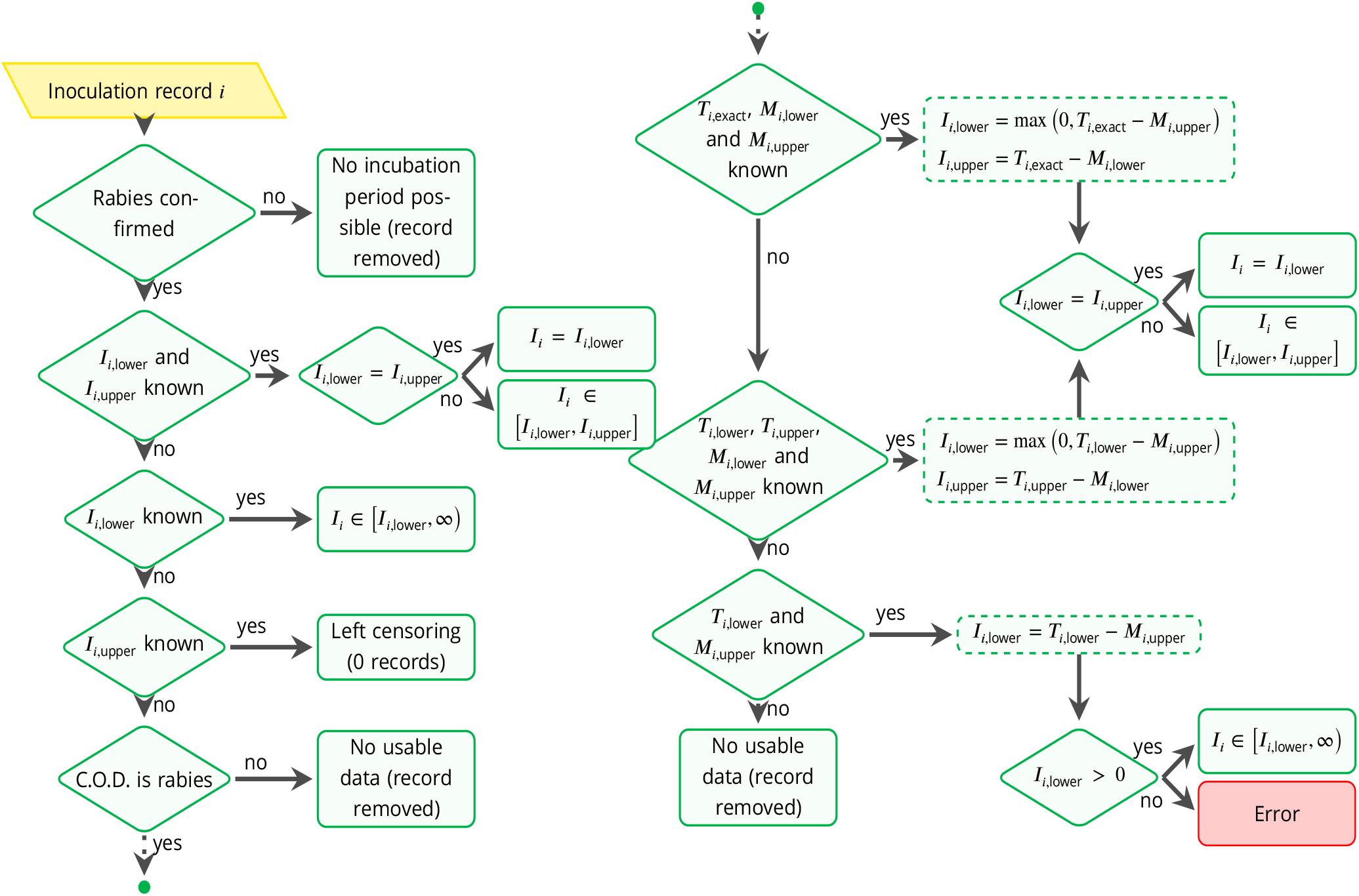
Overview of the algorithm used to consolidate data on incubation periods.

**Supplementary figure S 13:**
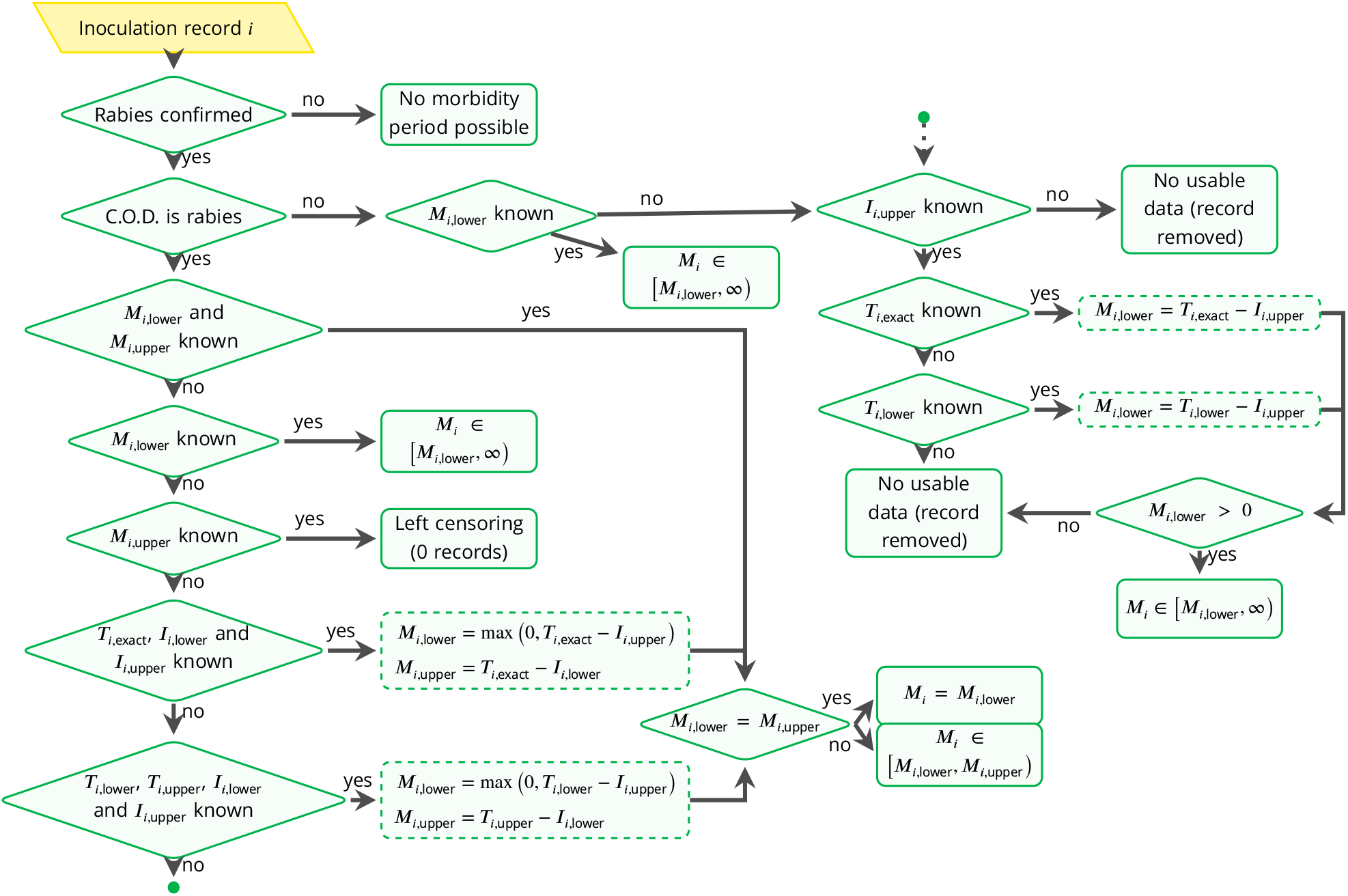
Overview of the algorithm used to consolidate data on clinical periods.

## Footnotes

1 www.itis.gov

## Notes

### Competing Interest Statement

The authors have declared no competing interest.

